# Munc13 binds and recruits SNAP25 to chaperone SNARE complex assembly

**DOI:** 10.1101/2020.08.19.257261

**Authors:** R Venkat Kalyana Sundaram, Huaizhou Jin, Feng Li, Tong Shu, Jeff Coleman, Jie Yang, Frederic Pincet, Yongli Zhang, Shyam S. Krishnakumar, James E. Rothman

## Abstract

Synaptic vesicle fusion is mediated by membrane-bridging complexes formed by SNARE proteins - VAMP2 on the vesicle and Syntaxin-1/SNAP25 on the pre-synaptic membrane. Accumulating evidence suggest that chaperones Munc18-1 and Munc13-1 co-operatively catalyze SNARE assembly via an intermediate ‘template’ complex containing Syntaxin-1 and VAMP2. How SNAP25 is chaperoned into this nascent complex remains a mystery. Here we report that Munc13-1 recruits SNAP25 to initiate the ternary SNARE complex assembly by direct binding, as judged by bulk FRET spectroscopy and single-molecule optical tweezer studies. Detailed structure-function analyses show that the binding is mediated by the Munc13-1 MUN domain and is specific for the SNAP25 ‘linker’ region that connects the two SNARE motifs. Consequently, freely diffusing SNAP25 molecules on phospholipid bilayers are concentrated and presumably bound in ~1:1 stoichiometry by the self-assembled Munc13-1 nanoclusters. Our data suggests that Munc13-1’s capacity to bind all three synaptic SNARE proteins likely underlie its chaperone function.

---

Neuronal communication involves the controlled release of neurotransmitters stored in synaptic vesicles (SV) into the neuronal synapse (Jahn and Fasshauer 2012; Rizo and Xu 2015; Sudhof 2013; Sudhof and Rothman 2009). This process is tightly regulated to ensure that the message is timely and precise (Sudhof 2013; Sudhof and Rothman 2009). SV fusion is catalyzed by the synaptic SNARE (soluble N-ethyl maleimide sensitive factor attachment protein) proteins - VAMP2 on the vesicle membrane (v-SNARE) and Syntaxin-1 and SNAP25 on the plasma membrane (t-SNAREs) (Sollner et al. 1993; Weber et al. 1998). When the vesicle approaches the plasma membrane (PM), the helical SNARE motifs of the cognate v- and t-SNAREs constitutively assemble into a ternary complex that initially bridges and ultimately fuses the two membranes (Sutton et al. 1998; Sollner et al. 1993; Weber et al. 1998). The SNARE complex nucleates into a four-helix bundle at its membrane distal end and progressively assembles (“zippers”) towards the membranes, exerting a potent force that ultimately drives the membranes together to fuse into one (Gao et al. 2012; Weber et al. 1998; Li et al. 2007). Though efficient, under *in vitro* conditions, the fusion process is artificially slow with its rate being limited by the intrinsic rate of nucleation of SNARE complex (Lai et al. 2017; Weninger et al. 2003; Pobbati et al. 2006). *In vivo*, nucleation is greatly accelerated by two co-operating, specialized molecular chaperones, Munc13 and Munc18 (Shen et al. 2007; Sudhof 2013; Sudhof and Rothman 2009; Ma et al. 2013). How precisely these chaperones and their homologues achieve this, be it in the highly specialized context of neurotransmission or more broadly for vesicle fusion driving intracellular protein transport, is a major open question despite important insights that have been gleaned.

Structural studies indicate that Munc18-1 binds and stabilizes monomeric Syntaxin-1 in a ‘closed’ (auto-inhibited) conformation (Dulubova et al. 2007; Misura et al. 2000). In doing so, it prevents its assembly into SNARE complexes or engagement into off-pathway reactions (Dulubova et al. 2007; Misura et al. 2000; Shen et al. 2007). This interaction is associated with Munc18-1 chaperone function and is critical for trafficking of Syntaxin-1 from endoplasmic reticulum (ER) to the PM (Shi et al. 2011; Rowe et al. 2001; Arunachalam et al. 2008). Munc18-1 has also been shown to promote SNARE complex assembly and to accelerate the resultant vesicular fusion (Shen et al. 2010; Shen et al. 2007). This activator function is attributed to Munc18-1’s ability to bind and sequester partially-assembled SNARE complexes, termed SNAREpins (Shen et al. 2007). Indeed, recent structural and biophysical studies suggest that Munc18-1 likely serves as a template for the formation of an early SNARE assembly intermediate (termed a ‘template complex’) wherein Syntaxin-1 and VAMP2 are properly oriented and aligned for nucleation into a four helix bundle (Baker et al. 2015; Shu et al. 2020; Jiao et al. 2018; Sitarska et al. 2017). Thus, the dual-binding modes of Munc18-1 to monomeric Syntaxin-1 and SNARE complexes allow it to both negatively-regulate and positively-assist SNARE complex assembly.

A key function of the other dedicated chaperone, Munc13-1, is to catalyze the overall process of nucleation by “opening” the closed Syntaxin-1/Munc18 complex to enable it to rapidly assemble with VAMP2 to form the template complex (Betz et al. 1997; Yang et al. 2015; Basu et al. 2005; Ma et al. 2011; Ma et al. 2013). This catalytic function has been mapped to the large central module within Munc13 called the MUN domain (Yang et al. 2015; Basu et al. 2005), but the precise mechanism is not known. Independent of Munc18-1, Munc13-1 has been shown to promote the proper alignment of Syntaxin-1 and VAMP2 (Lai et al. 2017) thereby accelerating and stabilizing the Munc18/Syntaxin-1/VAMP2 template complex (Shu et al. 2020). The mechanism involved was recently clarified with the findings that Munc13-1 independently binds both VAMP2 (Wang et al. 2019) and Syntaxin-1 (Wang et al. 2017; Ma et al. 2011) at distinct sites of its MUN domain potentially enabling it to simultaneously interact with and orient these two SNAREs in molecular proximity to Munc18-1.

Munc13-1 is a large (~20 nm) multi-domain, multi-functional protein (Xu et al. 2017). Independent of chaperoning SNAREpin assembly, Munc13-1 plays a key role in local tethering of SV at dedicated active zones of the pre-synaptic PM (Sudhof 2013; Rizo and Sudhof 2012; Camacho et al. 2017). Local tethering of SV to the PM, occurs prior to nucleation of SNAREpins, and is required to form the pool of release ready vesicles (Quade et al. 2019). It has been proposed that local tethering occurs when Munc13-1 itself forms molecular bridges linking the SV and PM through interactions involving the C_1_-C_2_B region and the C_2_C domain on opposite ends of the MUN domain (Quade et al. 2019; Liu et al. 2016; Shin et al. 2010; Basu et al. 2007). Specifically, the C_1_ and C_2_B domains bind the PM through interactions with their physiological ligands diacylglycerol (DAG) and phosphatidylinositol 4,5-bisphosphate (PIP2) respectively, while the C-terminal C_2_C domain interacts with the SV membrane, at least in part via the acidic phospholipid, phosphatidylserine (PS) (Liu et al. 2016; Quade et al. 2019; Shin et al. 2010; Michelassi et al. 2017; Kabachinski et al. 2014).

Overall, Munc18-1 and Munc13-1 co-operatively catalyze efficient SNAREpin nucleation via a template complex involving choreographed binding of Syntaxin-1 and VAMP-2. Of course, SNAREpins cannot nucleate without two of their four helices. Remarkably, especially for such a heavily investigated subject, how SNAP25 is chaperoned into nascent SNAREpins remains a mystery. It is known that VAMP2-containing SVs dock to PIP2-enriched regions on the PM, guided by Munc13-1 and SV-associated protein, Synaptotagmin (Park et al. 2015; van den Bogaart et al. 2012). Syntaxin-1 is sequestered into these PIP2 microdomains, mediated by electrostatic interactions involving the Syntaxin-1 juxta-membrane region (Honigmann et al. 2013). However, SNAP25, is highly dispersed on the PM surface and in fact, largely excluded from the PIP2 clusters under resting conditions (Chamberlain et al. 2001; Salaun et al. 2005). Exactly how SNAP25 is recruited to the site of vesicle docking to initiate SNARE complex assembly remains enigmatic.

Recently, a single-molecule force spectroscopy study revealed that the Munc13-1 MUN domain enhances the probability of SNAP25 binding to the Syntaxin-1/VAMP2/Munc18 template complex and subsequently complete assembly of the ternary SNARE complex (Shu et al. 2020). This suggested that Munc13-1 may play a role in recruiting SNAP25 to initiate SNARE complex formation. Here we test and confirm this hypothesis. We find that the Munc13-1 MUN domain directly and specifically binds the SNAP25 linker region and this interaction forms the molecular basis of Munc13-1 function in stimulating SNARE complex assembly both on its own and in conjunction with Munc18-1.

## RESULTS

### Munc13-1 MUN domain directly binds SNAP25 in solution

To test if there is direct molecular interaction between Munc13-1 and SNAP25, we employed Microscale Thermophoresis (MST), a highly sensitive technique that allows one to quantify molecular interactions with limited amounts of protein under native buffer conditions (Duhr and Braun 2006; Wienken et al. 2010). Informed by earlier reports (Liu et al. 2016; Quade et al. 2019; Wang et al. 2019; Yang et al. 2015; Wang et al. 2017), we focused on the conserved C-terminal portion of Munc13-1 consisting of the contiguous C_1_-C_2_B-MUN-C_2_C domains (Munc13-1 residues 529-1735; referred to as Munc13_L_), except for residues 1408-1452 within a non-conserved loop in the MUN domain, which was deleted and replaced with EF residues to minimize dimerization/oligomerization (Basu et al. 2005; Ma et al. 2011). To facilitate the MST analysis, we introduced a C-terminal Halo tag to the Munc13_L_ fragment to enable covalent attachment of a fluorescent tag. Titration of full-length wild-type SNAP25 (residues 1-206, SNAP25^WT^) into AlexaFluor 660 labeled Munc13_L_ yielded a classical single-site dose-response curve with a dissociation constant (K_d_) of 26 ± 2 μM (Figure 1A). This indicated that that Munc13-1 directly binds SNAP25, albeit with a modest affinity under soluble conditions. This binding affinity is comparable to that reported (25-100 μM) for specific binding of the MUN domain to Syntaxin-1, VAMP2, and the fully assembled SNARE complex when these proteins are also free in solution (Wang et al. 2017; Wang et al. 2019; Ma et al. 2011; Lai et al. 2017). Therefore, the novel binding we observe between Munc13-1 and SNAP25 is similarly likely to be physiologically relevant at the high local concentrations that exist in active zones.

**Figure 1.**
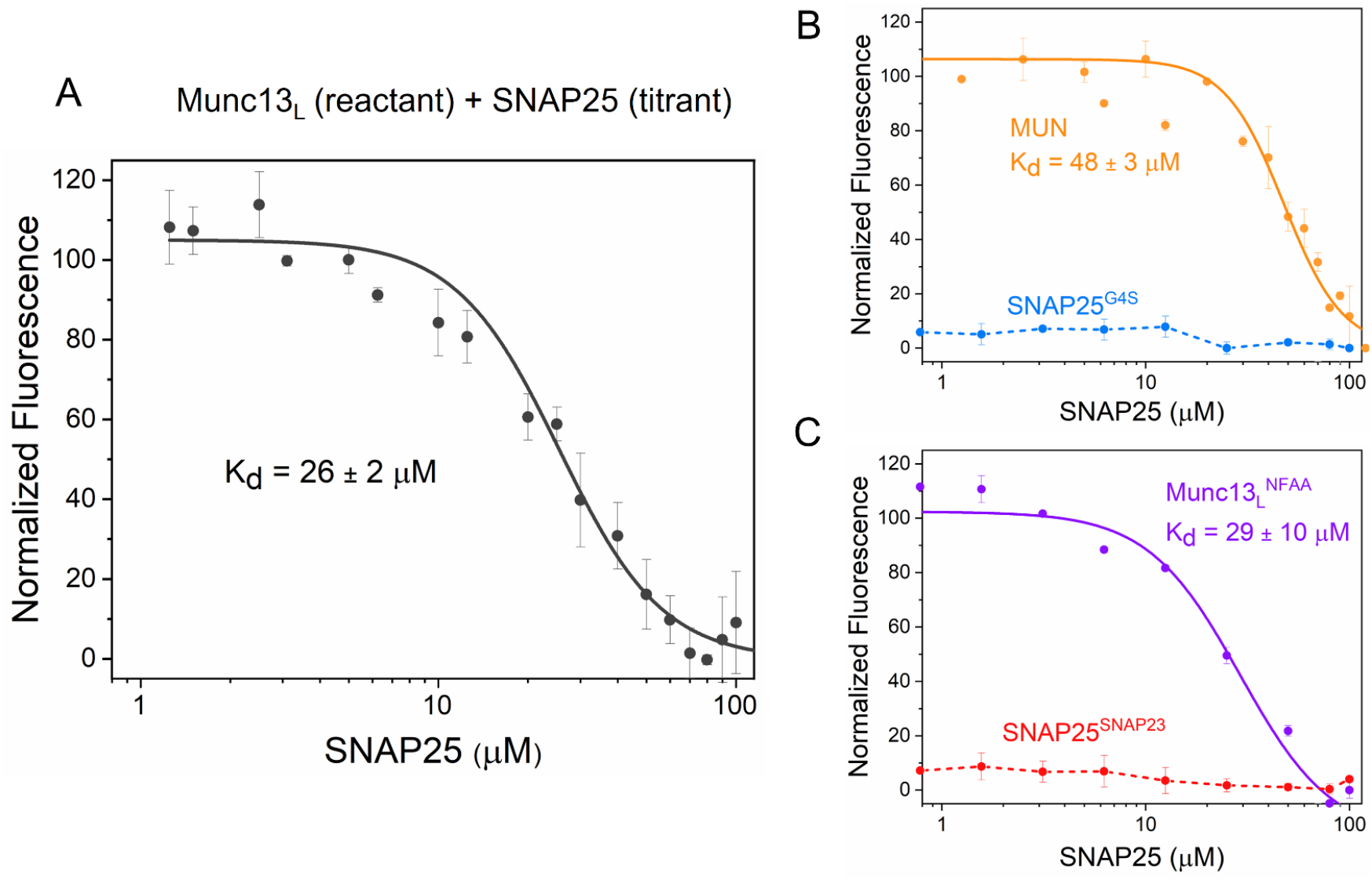
Dissecting Munc13-1 interaction with SNAP25 using Microscale Thermophoresis (MST) analysis. **(A)** Titration of full-length SNAP25 into AlexaFluor 660 labeled Munc13_L_ (C1-C2B-MUN-C2C domains) produced a concentration-dependent change in the MST signal, yielding an apparent dissociation constant (K_d_) = 26 ± 2 μM implying that Munc13_L_ directly binds to SNAP25. **(B)** MUN domain (orange curve) alone is sufficient to bind SNAP25 albeit with slightly reduced affinity (K_d_ = 48 ± 3 μM) under similar experimental conditions. However, replacing the SNAP25 linker domain that connects that SNARE helical motifs with a GGGGS sequence (SNAP25^G4S^) completely abrogates Munc13-SNAP25 interaction (blue curve). This suggests that the Munc13-1 MUN domain likely binds to the SNAP25 linker region. **(C)** MUN-SNAP25 linker region interaction is highly specific as replacing the SNAP25 linker region with closely associated SNAP23 linker (SNAP25^SNAP23^) also disrupts the binding (red curve). On the other hand, Syntaxin non-binding Munc13-1 mutant (Munc13_L_^NFAA^) is capable of binding SNAP25 (purple curve) indicating that the SNAP25 binding site is unique and different from the Syntaxin-1 binding site. Average and standard deviations from a minimum of three independent experiments are shown.

We then sought to identify the regions within Munc13-1 and SNAP25 responsible for their binding. The central MUN domain of Munc13-1 contains the distinct binding sites for both VAMP2 and Syntaxin-1, each of which is required for its chaperone function (Wang et al. 2017; Wang et al. 2019). So, we expressed and purified the isolated MUN domain (residues 859-1531 Δ1408-1452, EF), and labeled it with AlexaFluor 660 at a C-terminal Halo tag. MST analysis showed that the MUN domain alone can bind SNAP25^WT^ with an apparent K_d_ of 48 ± 3 μM (Figure 1B). This indicated that the MUN domain serves as the primary binding site for SNAP25 and that the flanking C_1_-C_2_B and C_2_C portions might weakly augment the primary MUN-SNAP25 binding site, be it directly or indirectly.

### Binding SNAP25 involves the linker region between its two SNARE motifs

Since the isolated MUN domain promotes the nucleation of SNARE complexes (Basu et al. 2005), we reasoned that the Munc13-1 binding site on SNAP25 is likely outside of the SNARE helices, as these would need to be free to assemble with the SNARE motifs of VAMP2 and Syntaxin-1. This logic would make the 60 residue ‘linker’ region that concatenates the SNARE helical motifs the best candidate for productive binding by Munc13-1. Therefore, we designed and tested a SNAP25 mutant (SNAP25^G4S^) in which the linker region was swapped with an inert, flexible, non-specific sequence (GGGGS repeats) yet maintaining the overall length. Even though SNAP25^G4S^ successfully replaced SNAP25^WT^ in a bulk vesicle fusion assay (Figure 1 Supplement 1) and has been shown to assemble in SNARE complex (Shaaban et al. 2019), it failed to bind to Munc13_L_ even at the highest SNAP25^G4S^ concentration (100 μM) tested (Figure 1B). This suggests that the MUN domain likely binds to the SNAP25 linker region.

### SNAP25 binding to Munc13-1 is specific and unique

To assess the specificity of binding, we replaced the SNAP25 linker region with that of its close homologue, SNAP23, producing a chimeric protein termed SNAP25^SNAP23^. SNAP23 is structurally and functionally similar to SNAP25 and is expressed in all tissues (Kunii et al. 2016; Ravichandran et al. 1996; Wang et al. 1997), where it associates with Syntaxin-1 and VAMP2 and functions as part of the ubiquitous cellular fusion machinery (Kunii et al. 2016). Notably, the linker region of SNAP23 (compared with SNAP25) is markedly less conserved (~45% identical) than either SNARE motif (~70%). Indeed, SNAP25^SNAP23^ failed to bind to Munc13_L_ (Figure 1C), confirming that Munc13-1 binding to SNAP-25 is isoform specific and that specificity resides in the linker region.

Munc13-1 has been shown to bind and activate Syntaxin-1 via a hydrophobic pocket near the middle of the MUN domain (Wang et al. 2017). To assess if SNAP25 binding overlaps with the Syntaxin site, we tested a previously described mutations in Munc13-1 (N1128A/F1131A; Munc13_L_^NF^) that disrupts binding to Syntaxin-1 (Wang et al. 2017). These mutations had no effect and Munc13_L_^NF^ bound SNAP25 with a K_d_ of 29 ± 10 μM (Figure 1C, purple). This indicates that the SNAP25 binding site within the MUN domain is distinct from the Syntaxin-1 binding pocket.

### Munc13-1 binds SNAP25 on lipid membrane surface at 1:1 stoichiometry

SNAP25 and Munc13 are both naturally concentrated within active zones of the pre-synaptic plasma membrane, where they are restricted to a common surface by distinct mechanisms. SNAP25 is anchored by the direct insertion of its multiple palmitoylated cysteines in the linker region (Greaves et al. 2009; Nagy et al. 2008; Shaaban et al. 2019). As mentioned, Munc13-1 is anchored by binding to lipid headgroups by C_1_-C_2_B domains (Liu et al. 2016; Shin et al. 2010; Basu et al. 2007). We thus investigated the SNAP25-Munc13 interaction on a lipid bilayer surface. We anticipated that restriction of these two proteins to a two dimensional surface would increase the association rate, resulting overall in markedly greater stability of the Munc13-SNAP25 complexes thereby enabling them to be imaged and quantified on the supported bilayers using total internal reflection fluorescence (TIRF) microscopy.

We created a supported lipid bilayer with physiologically relevant lipid composition (73% PC, 15% PS, 3% PIP2, 2% DAG) by bursting pre-formed vesicles onto a glass coverslip. We had anticipated that it would be necessary to use palmitoylated SNAP25 in this experiment, but instead observed that even non-palmitoylated SNAP25, added at low (150 nM) concentrations in solution, bound to the bilayer. This is likely due to the amphipathic nature of two SNARE motifs of SNAP25 (Sutton et al. 1998) as it is well-established that amphipathic peptides associate strongly to phospholipid surfaces as their hydrophobic side chains could insert into the lipid bilayers (Wagle et al. 2019; Gimenez-Andres et al. 2018). On its own, freely-diffusing AlexaFluor 555-labeled SNAP25 molecules thus bound were monomeric and uniformly distributed on the lipid bilayer surface (Figure 2A, Figure 2 Supplement 1). When Munc13_L_ was present, SNAP25 was no longer monodisperse, but rather located in numerous distinct clusters of variable intensity (Figure 2A) and the clustering efficiency directly correlated to the concentration of Munc13_L_ (Figure 2A).

**Figure 2.**
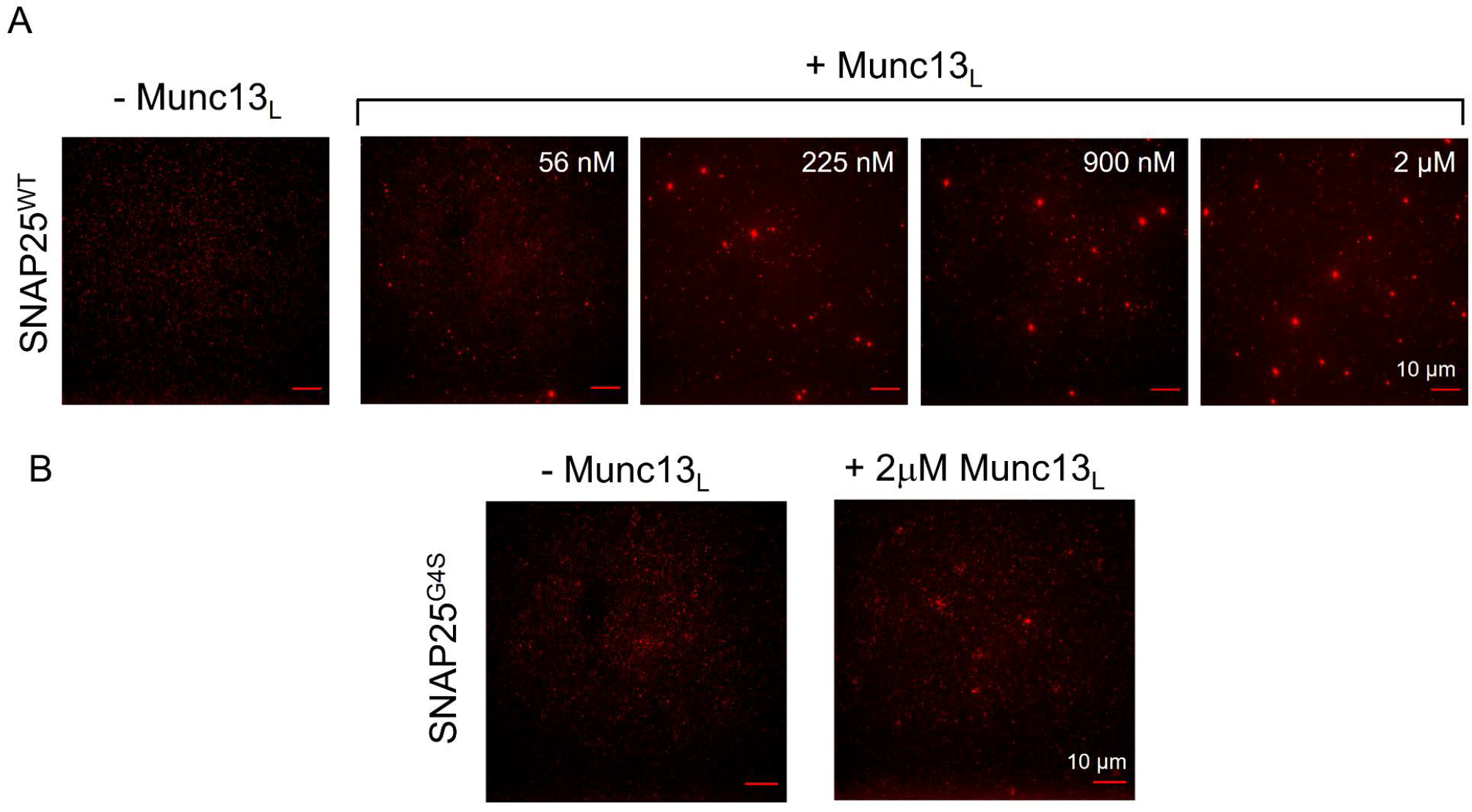
Munc13-1 binds and clusters SNAP25 on lipid bilayer surface. **(A)** Distribution of AlexaFluor 488 labeled SNAP25^WT^ (~150 nM) on supported lipid bilayer under physiological lipid and buffer composition in the absence or presence of Munc13_L_ was visualized using TIRF microscopy. Munc13_L_ induces clustering of SNAP25^WT^ molecules and the number/size of the SNAP25 clusters directly correlates to concentration of Munc13_L_ added. **(B)** Munc13_L_ fails to cluster SNAP25 linker domain mutant (SNAP25^G4S^) even at the highest concentration (2 μM) tested. This suggests that MUN-SNAP25 linker domain interaction is feasible on lipid bilayers surfaces and results in clustering of otherwise diffusely distributed SNAP25 molecules. Representative fluorescent images are shown.

Considering that Munc13-1 has been shown to self-associate into clusters on lipid bilayers under both *in vitro* and physiological conditions (Sakamoto et al. 2018; Li et al. 2020), we reasoned that SNAP25 puncta are a consequence of its association with Munc13-1 clusters. To confirm, we co-localized AlexaFluor555-labeled SNAP25 with AlexaFluor488-labeled Munc13_L_ using two color TIRF microscopy (Figure 3A). As expected, Munc13_L_ on its own formed clusters on lipid bilayer surface (Figure 3A) and SNAP25 increasingly co-localized with these clusters as the concentration of Munc13_L_ was increased. When Munc13_L_ was added at 50 nM bulk concentration the Manders’ coefficient (SNAP25:Munc13_L_) was 0.2 ± 0.07, which increased to 0.6 ± 0.05 at 1 μM Munc13_L_, consistent with a concentration-dependent binding process. As a control, we employed membrane-anchored (palmitoylated) SNAP25 and obtained similar results (Figure 3 Supplement 1).

**Figure 3.**
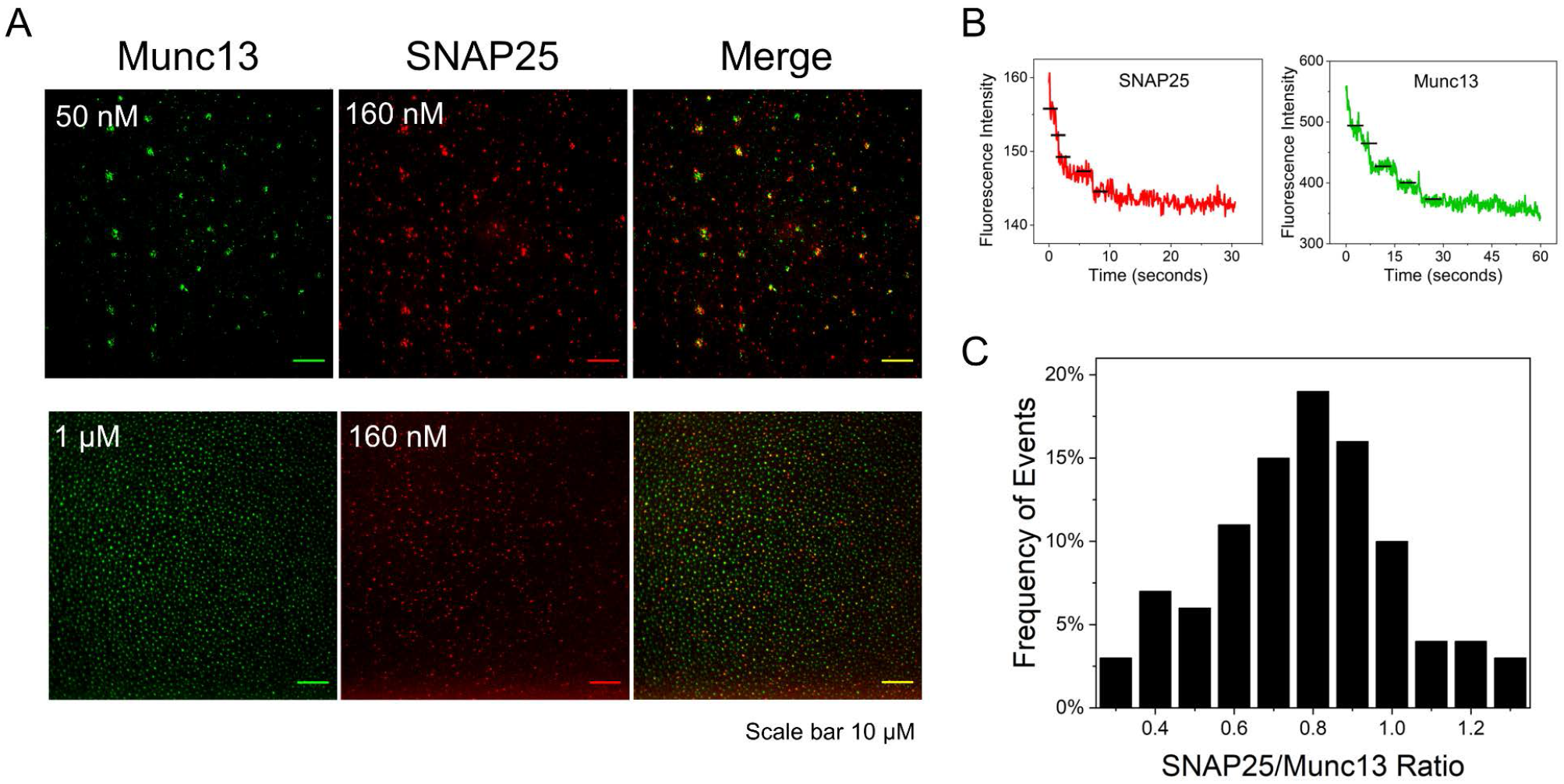
Munc13-1 co-clusters with SNAP25 at 1:1 stoichiometry. **(A)** Representative fluorescence images showing that AlexaFluor 555 labeled SNAP25^WT^ co-localizes with clusters of AlexaFluor 488 labeled Munc13_L_ on lipid bilayer surface and the extent of co-localization depends on the concentration of Munc13_L_ added. SNAP25 (160 nM) shows limited co-localization (Manders’ coefficient ~ 0.2) at 50 nM of Munc13_L_ but has high overlap (Manders’ coefficient ~ 0.7) at 1μM of Munc13_L_. **(B)** Stoichiometry of SNAP25 and Munc13_L_ in the co-clusters under low Munc13_L_ concentration was estimated using step-bleaching analysis. Representative step-bleaching traces for SNAP25 (red) and Munc13_L_ (green) are shown and the ratio of SNAP25:Munc13_L_ quantified from ~70 co-clusters is shown as the histogram. Under these conditions, each cluster contains ~5-7 molecules at almost 1:1 ratio of Munc13_L_:SNAP25. This indicates that molecular interaction between the MUN domain and the SNAP25 linker region is likely responsible for the observed co-clustering of Munc13-1 and SNAP25.

To establish the specificity of this binding, we employed the linker mutant, SNAP25^G4S^. As expected, no clustering of SNAP25^G4S^ was observed, even at the highest concentration (2 μM) of Munc13_L_ (Figure 2B). This suggests that similar to the soluble conditions, SNAP25 binding to Munc13_L_ nanoclusters on membranes involves the SNAP25 linker region. Finally, to establish the stoichiometry of binding of SNAP25 to the Munc13_L_ nanoclusters, we performed two color photo-bleaching experiments (Fig. 3B). Reliable identification of single steps by this method can only be made when relatively small (<10) numbers of molecules are present.

Therefore, we performed these experiments at a low Munc13_L_ concentration (50 nM) designed to limits the copy number of Munc13_L_ in the nanoclusters. Among these co-clusters, independent of their size, we observed a broad distribution of the apparent mole ratio of SNAP25 to Munc13_L_ centering on ~0.8 mole/mole (Figure 3B). Considering the high and similar efficiency of labeling of both SNAP25 and Munc13_L_, the ratio of fluorescence likely reflects the actual mole ratio. This suggests that Munc13-1 binds SNAP25 at approximately 1:1 stoichiometry.

### Linker-specific binding by MUN is necessary for SNAP25 to enter the SNARE complex

We then examined the potential functional relevance of the MUN-SNAP25 linker domain interaction to Munc13-1’s established and independent role (Lai et al. 2017; Liu et al. 2016; Shu et al. 2020) in accelerating nucleation of SNAREpins. We employed a fluorescence resonance energy transfer (FRET) assay with fluorescent probes (AlexaFluor 488 & AlexaFluor 555) introduced at the N-termini of SNAP25 (residue Q20) and VAMP2 (residue S28) respectively to directly track the effect of the MUN domain on the initiation of SNARE complex assembly.

In the absence of the MUN domain, SNAP25^WT^ spontaneously assembled into a complex with VAMP2 and unlabeled Syntaxin-1 over the course of an hour as evident from the gradual increase of the acceptor fluorescence intensity (Figure 4A). We did not detect FRET in control experiments when Syntaxin-1 was omitted confirming that the observed FRET signal corresponds to the formation of a ternary SNARE complex (Figure 4A). In line with the bulk-fusion assays (Figure 1 Supplement 1) and previous reports (Nagy et al. 2008; Shaaban et al. 2019), mutation of the linker region had very little effect on the spontaneous rate of SNAP25 assembly into SNARE complexes (Figure 4A), with comparable increase in acceptor (FRET) signal with SNAP25^G4S^ mutant (Figure 4A). The MUN domain strongly stimulated the rate of association of SNAP25^WT^ with the cognate SNAREs (~3-fold enhancement of FRET signal after 60 min) but had no effect on the rate of assembly of SNAP25^G4S^ mutant (Figures 4A and 4B). This data forcefully suggests that binding of SNAP25 (via its linker) is required for Munc13-1 to chaperone SNARE complex assembly.

**Figure 4.**
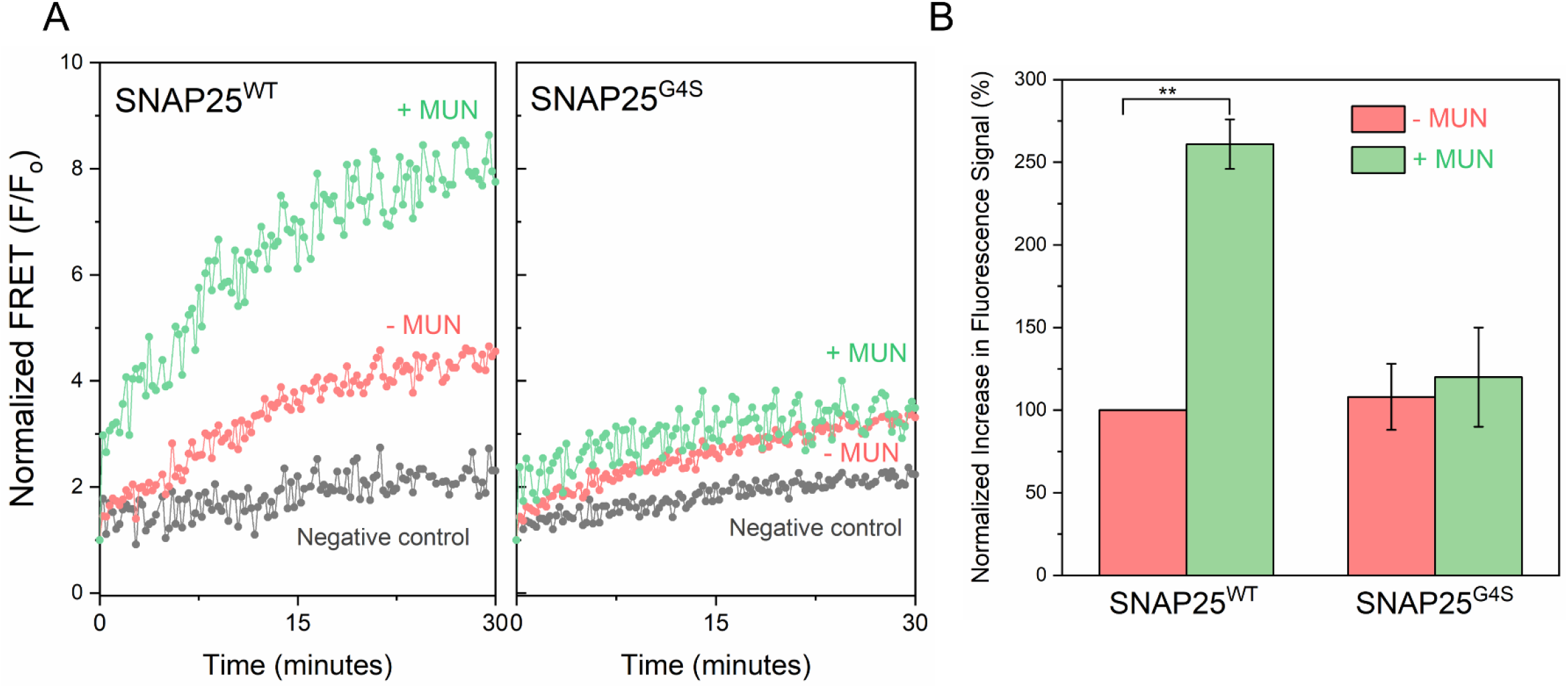
Munc13 interaction with SNAP25 is essential for its chaperone function. FRET between AlexaFluor 488 labeled SNAP25 and AlexaFluor 555 labeled soluble VAMP2 introduced in the N-terminus (SNAP25 residue 20 & VAMP2 residue 28) shows that in the presence of cytoplasmic Syntaxin-1, the MUN domain activates the initial engagement of the SNARE complex. Negative control wherein Syntaxin-1 was left out confirms the observed FRET signal corresponds to formation of the ternary SNARE complex. The stimulatory effect of MUN domain was not observed with SNAP25 linker mutant (SNAP25^G4S^) indicating that the MUN-SNAP25 linker domain interaction is crucial for Munc13-1 ability to promote SNARE complex formation. Representative fluorescence traces for first 30 min of the reaction (left) and end-point quantification at 60 min (right) from two independent trials are shown. Statistical significance was determined using student t-test (** p<0.005).

### Linker-dependent binding by MUN recruits SNAP25 to the template complex

To independently test this conclusion in the fuller context of Munc13-1’s co-operation with Munc18-1 in chaperoning SNARE complex assembly, we employed an established single molecule assay involving optical tweezers (Figure 5A) (Jiao et al. 2017; Rebane et al. 2016; Jiao et al. 2018; Shu et al. 2020). Here, Syntaxin-1 and VAMP2 were cross-linked at their N-termini (between Syntaxin R198C and VAMP2 N29C) via a disulfide bond and pulled from their C-termini via two DNA handles attached to two optically trapped polystyrene beads (Shu et al. 2020; Jiao et al. 2018). We first pulled a single SNARE complex in the presence of Munc18-1 and SNAP25^WT^ (0.5 μM & 0.6 μM respectively added into the solution). The resultant force-extension curve (FEC) indicated that a single SNARE complex unfolds in a characteristic stepwise manner among different states when pulled (Figure 5B, FEC #1 & #2 grey curves with different regions indicated by their associated state numbers). We observed reversible C-terminal domain (CTD) transitions at about 15 pN (between state 1 and state 2, regions marked by gray ovals), followed by an irreversible unfolding of the N-terminal domain (NTD) at ~16-17 pN (from state 2 to state 3, extension jumps marked by gray arrows), and finally irreversible unfolding of the Syntaxin-SNAP25 (t-SNARE) complex (marked by green arrows) with the attendant dissociation of SNAP-25 (state 5).

**Figure 5:**
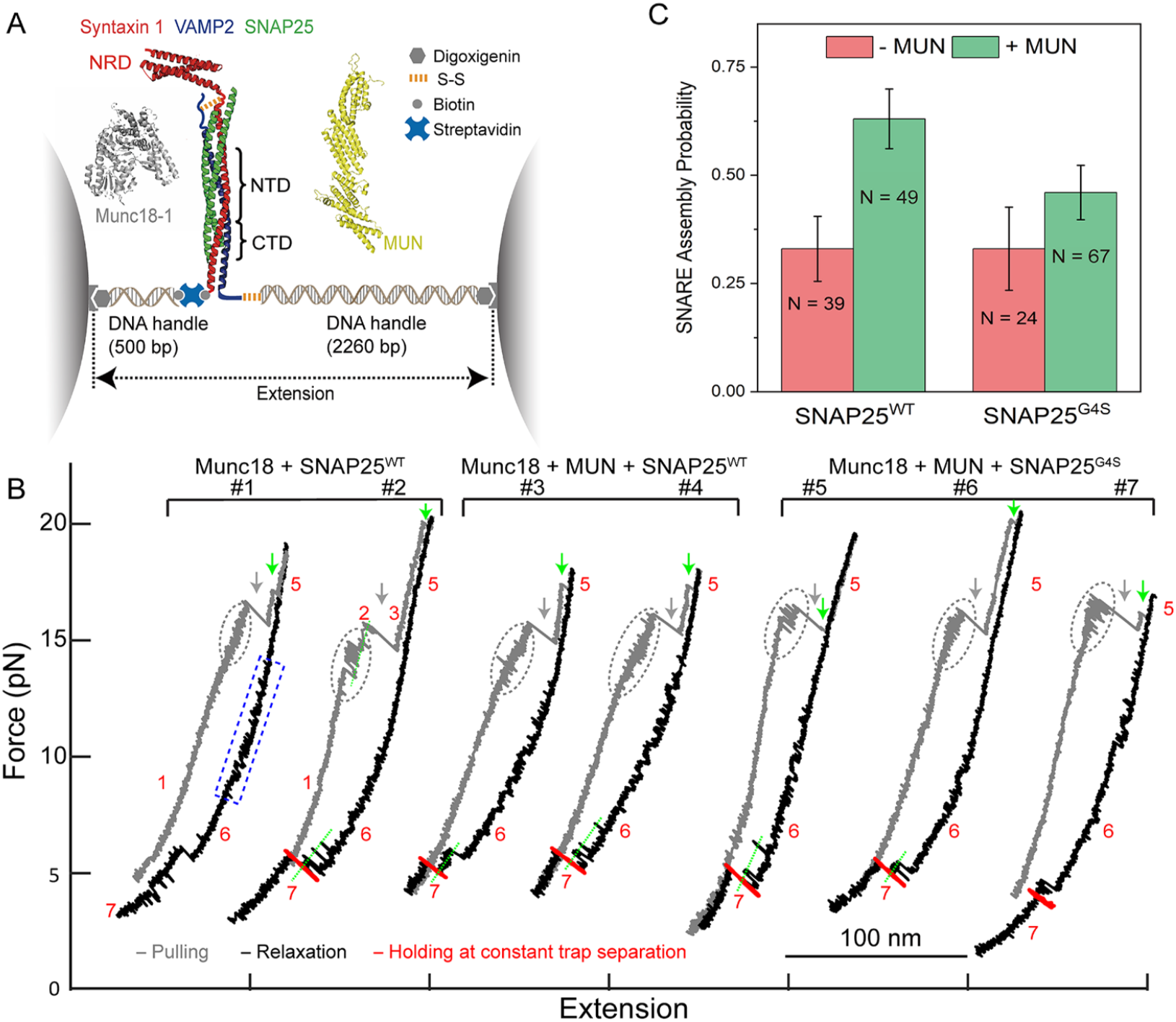
Munc13-1 recruits SNAP25 to Munc18-1/Syntaxin-1/VAMP2 template complex. **(A)** Experimental setup for optical tweezers. A single SNARE complex was pulled from the C termini of Syntaxin 1A (red) and VAMP2 (blue) via two DNA handles attached to two optically trapped polystyrene beads. The N-termini of Syntaxin 1A and VAMP2 were cross-linked via a disulfide bond. Munc18-1 (gray) and the MUN domain of Munc13-1 (yellow) were added in the solution. **(B)** Representative force extension curves (FECs) obtained for SNARE complexes assembled with SNAP25^WT^ or SNAP25^G4S^ in the presence of Munc18-1 and MUN domain as indicated. The Syntaxin–VAMPs conjugate was pulled (grey FECs) or relaxed (black FECs) by changing the separation between two optical traps at a speed of 10 nm/s or held at constant mean force around 5 pN (red FECs). The molecular states associated with different FEC regions are indicated by the corresponding state numbers (see Figure 2D in ref. (Jiao et al. 2018) for their molecular diagrams) **(C)** Probability of full SNARE complex assembly from the Munc18-1/Syntaxin-1/VAMP2 template complex observed within 100s at 5 pN constant mean force. MUN domain ability to stimulate SNARE complex assembly is significantly diminished with the SNAP25 linker domain mutant (SNAP25^G4S^) as compared to SNAP25^WT^. The N value refers to the total number of trials and average and standard error of the means are shown.

Relaxing the force applied to the SNARE proteins then revealed two reversible transitions along the relaxation curves (Figure 5B FEC #1 black curve). The first transition from 14 to 6 pN (marked by blue rectangle) was caused by the folding of partially closed Syntaxin-1 (state 6) and the lower force transition at ~3-6 pN resulted from the formation of Munc18-1/Syntaxin-1/VAMP2 template complex (state 7).

To detect the ternary SNARE complex assembly, we held the template complex at a constant mean force around 5 pN for 100 seconds to await SNAP25^WT^ binding (Figure 5B FEC#2 red curve). Indeed, in some cases, SNAP25^WT^ from the solution was able to bind to the template complex resulting in reassembly of the full SNARE complex (state 1). This reflects the intrinsic rate of productive association of free SNAP25 with the template complex.

To test whether the rate of productive association of SNAP25 to the template complex is accelerated by Munc13-1, we included the MUN domain (1 μM) in addition to Munc18-1 and SNAP25^WT^ in solution (Figure 5B FEC #3 & #4). Indeed, the MUN domain stabilized the template complex and promoted the rate of re-assembly of the SNARE complex from the template complex. Statistically, the MUN domain nearly doubled the probability of SNARE assembly from the template complex (Figure 5C). All of these observations are consistent with our previous reports (Shu et al. 2020; Jiao et al. 2018).

To test whether binding of SNAP25 to Munc13-1 via the SNAP25 linker is required for Munc13-1 to chaperone SNAP25 in this context, we repeated the above experiment with linker domain mutant, SNAP25^G4S^. With Munc18-1 only, the probability of SNARE complex formation with the SNAP25^G4S^ was similar to that observed with SNAP25^WT^ (Figure 5C). This further confirmed the alteration of the liker domain does not affect the inherent ability to SNAP25 helices to bind its cognate SNARE partners. Addition of the MUN domain along with Munc18-1 (Figure 5B, FEC #5, #6, #7) stabilized the template complex (Shu et al. 2020) but did not appreciably increase the probability of the SNAP25^G4S^ to complete SNARE assembly as compared to SNAP25^WT^ (Figure 5C). This strongly indicates that MUN-SNAP25 binding (via the linker domain) contributes to the enhanced assembly of the four-helix bundle from the Munc18-1/Syntaxin-1/VAMP2 template complex.

## DISCUSSION

Altogether, we have identified a novel, functionally relevant interaction between the Munc13-1 MUN domain and the SNAP25 linker region. Our finding is supported by several previous studies (Dubuke et al. 2015; Sivaram et al. 2005; Nagy et al. 2008; Shaaban et al. 2019). For example, it has been demonstrated that yeast exocyst protein Sec6p (which is strikingly similar to the c-terminal half of Munc13-1 MUN domain) binds Sec9 (a SNAP25 homologue) via the linker region, and disrupting the Sec6p-Sec9 interaction results in significant defects in exocytosis and overall cell growth (Dubuke et al. 2015; Sivaram et al. 2005). Furthermore, the SNAP25 linker domain has been shown to play a crucial role in regulating vesicle priming and the Ca^2+^-regulated exocytosis in chromaffin cells (Nagy et al. 2008; Shaaban et al. 2019). Cellular analysis revealed that SNAP25 linker-domain mutant (similar to the SNAP25^G4S^ used in this study), which has an innate capacity to form a productive SNARE complex, fails to rescue secretion and significantly reduces the size of primed vesicle pools (Shaaban et al. 2019). This correlates to impairments in Munc13-1 chaperoned SNARE complex assembly observed in the *in vitro* FRET and optical-tweezer experiments (Figure 4 & 5).

This newly uncovered feature of Munc13-1 solves an enduring mystery – how is SNAP25 chaperoned into the SNARE complex – while being compatible with the several established roles of this remarkably multi-functional protein. We now see that Munc13-1 can potentially bind all three synaptic SNAREs simultaneously to sterically position them along with Munc18-1 to maximize the rate of their productive assembly into SNAREpins (Lai et al. 2017; Ma et al. 2013). It is well-established that the inherently cytoplasmic Munc13-1 translocates to DAG & PIP_2_-rich domains on the PM in response to neuronal activity and elevated intracellular Ca^2+^ concentration (Quade et al. 2019; Michelassi et al. 2017). On the PM, Munc13-1 initially functions as a tethering factor for local capture of SVs (Quade et al. 2019; Rizo and Sudhof 2012; Sudhof 2013; Li et al. 2020). At this stage, Munc13-1 is thought to adopt an *‘erect’* topology with the C_1_/C_2_B region anchored on the PM and extended (~ 15 nm) MUN domain reaching out to tether the SV via the C_2_C domain at its apex (Gipson et al. 2017; Quade et al. 2019; Xu et al. 2017). However, this *erect* topology seems incompatible with Munc13-1 role in catalyzing the nucleation of SNAREpins because in this extended configuration, Munc13-1 would prevent the membrane from coming closer and any nucleated SNAREpin would be inherently unstable when the involved bilayers are separated by more than ~ 10 nm (Wang et al. 2016; Li et al. 2007).

Munc13-1 is therefore predicted to change its orientation, ultimately to a *‘flat’* topology, in which the inter-membrane separation is reduced to ~5 nm (as observed in release-ready vesicles in the active zone) where SNAREpins can be meta-stably clamped at the approximately half-zippered state (Li et al. 2019; Hua and Charlton 1999; Prashad and Charlton 2014). This reorientation is likely driven by rearrangement of the C1-C2B domains on the PM (Michelassi et al. 2017; Liu et al. 2016), and may hypothetically occur via transitional states at intermediate separations that sterically favor the concerted reaction in which three SNAREs from two membranes and two molecular chaperones can favorably align. The precise molecular details of Munc13-1’s activation from a local tether to a SNARE chaperone remains to be determined. In this context, we cannot adjudicate whether Munc13-1 can sterically bind SNAP25 in the *erect* conformation or requires a conformational switch because the location of the binding site within the MUN domain is not yet known.

DAG/PIP_2_ binding has been shown to induce recruitment and clustering of Munc13-1 on the PM surface (Michelassi et al. 2017; Li et al. 2020; Sakamoto et al. 2018) and nanoclusters containing ~5-10 Munc13-1 molecules have been shown to be intimately associated with SV in the readily-releasable pool by super-resolution microscopy (Sakamoto et al. 2018; Reddy-Alla et al. 2017). Recently, we have shown that possibly similar nanoclusters form by self-assembly of Munc13-1 protomers (Li et al. 2020). Local multi-valency likely augments Munc13’s chaperone functions as it does vesicle capture (Quade et al. 2019; Li et al. 2020). It is easy to imagine that Munc13-1 clusters could help concentrate SNAP25 molecules at the vesicle docking site, similar to the co-cluster we observe (Figure 3) thus increasing the probability of productive SNARE complex formation. In summary, we address a long-standing question concerning the recruitment of SNAP25 to the vesicle docking site and in turn, highlight the pivotal role of Munc13-1 in orchestrating SNARE complex assembly.

## MATERIALS AND METHODS

### Materials

The cDNA constructs used in this study include full-length mouse His^6^-SNAP25b, residues 1–206 in a pET28 vector with a N-terminal Thrombin cut site; His^12^-Munc13_L_ (Munc13-1 residues 529-1735 with residues 1408–1452 replaced by the sequence EF) in a modified pCMV-AN6 vector which includes a PreScission cut site following the His^12^ tag, GST-MUN (residues 859-1407, Δ1408-1452, EF) cloned into a pGEX vector with a N-terminal Thrombin cut site (a kind gift from Dr. Josep Rizo, UT Southwestern). We used a QuikChange mutagenesis kit (Agilent Technologies, Santa Clara, CA) to generate the N1128A/F1131A in the Munc13_L_ background (Munc13_L_^NF^). SNAP25 linker domain mutants were synthesized (Genewiz Inc., Plainfield, NJ) with the SNAP25 residues 83-142 replaced with (GGGGS)_10_ repeats (SNAP25^G4S^) or SNAP23 residues 83-142 (referred to as SNAP25^SNAP23^) and cloned into the pET28 vector similar to the SNAP25^WT^ construct. For fluorescent labeling experiments, we introduced a Halo tag at the c-terminus of the Munc13_L_ and MUN constructs in the aforementioned vector background. We used a single-cysteine (S28C) version of SNAP25^WT^ (Krishnakumar et al. 2013) and SNAP25^G4S^ (synthesized from Genewiz Inc., Plainfield, NJ). Constructs used in the optical tweezer experiments (cytoplasmic domains of rat syntaxin 1A residues 1-265, R198C and VAMP2 residues 1-96, N29C) and the bulk FRET experiments (VAMP2 residues 1-96, S28C, SNAP25 residues 1-206, Q20C, Syntaxin 1 residues 1-258) have been described previously (Jiao et al. 2017; Shu et al. 2020). Lipids, 1,2-dioleoyl-sn-glycero-3-phosphocholine (DOPC), 1,2-dioleoyl-sn-glycero-3-phospho-L-serine) (DOPS), 1,2-dioleoyl-sn-glycero-3-phosphoethanolamine-N-[amino(polyethylene glycol)-2000] (DOPE-PEG2000), phosphatidylinositol 4, 5-bisphosphate (PIP2), diacylglycerol (DAG) and 1,2-dipalmitoyl-sn-glycero-3-phosphoethanolamine-N-(7-nitro-2-1,3 benzoxadiazol-4-yl) (NBD-PE) were purchased from Avanti Polar Lipids (Alabaster, AL). The polystyrene beads (2.1 μm in diameter) with anti-digoxigenin coating used in optical tweezer experiments were purchased from Spherotech (Green Oaks, IL).

### Protein production and purification

The MUN domain (with and without Halo tag) and all SNARE proteins were expressed and purified as described previously (Shu et al. 2020; Wang et al. 2016; Yang et al. 2015). Briefly, the proteins were expressed in Escherichia coli strain BL21(DE3) (Novagen, Darmstadt, Germany) and cells were lysed with a cell disruptor (Avestin, Ottawa, Canada) in lysis buffer containing 50 mM HEPES, pH 7.4, 400 mM KCl, 4% TritonX-100, 10% glycerol, 0.5 mM Tris (2-carboxyethyl) phosphine hydrochloride (TCEP) and 1 mM phenylmethylsulfonyl fluoride. Samples were clarified using a 45 Ti rotor (Beckman Coulter, Brea, CA) at 140,000 x g for 30 minutes and incubated with Ni-NTA agarose (Thermofisher, Waltham, MA) or Glutathione agarose (Thermofisher, Waltham, MA) for 4–16 h at 4°C. The resin was subsequently washed in the HEPES buffer, without TritonX-100 and supplemented with 50 mM Imidazole for the Ni-NTA purification. After the washes (three column volumes) the protein was cleaved of the resin with Thrombin for 4–16 h at 4°C. The protein was purified using gel filtration (Superdex75 or 200 Hi-load column, GE Healthcare, Chicago, IL). The peak fractions were pooled and concentrated using filters of appropriate cut-offs (EMD Millipore, Burlington, MA). NOTE: Syntaxin-1 construct used in optical tweezer studies was biotinylated *in situ* using previously described biotin ligase protocol (Jiao et al. 2017).

Munc13_L_ (with or without Halo-tag) was expressed in ExpiHEK-293 cell cultures using expifectamine as a transfection reagent (Thermofisher, Waltham, MA). Briefly, cells were passaged three times prior to transfection and were grown for 72 hours before being spun down and rinsed in ice cold PBS. The pellet was re-suspended in the HEPES lysis buffer and lysed using a Dounce homogenizer. The sample was clarified using at 140,000 x g for 30 minutes at 4°C and the supernatant incubated overnight with Ni-NTA beads, in the presence of DNAse 1, RNAse A and Benzonase to remove nucleotide contamination. The protein was further washed in the lysis buffer (without TritonX-100) before being cleaved with PreScission Protease for 2 hours at room temperature. The eluted proteins were further purified via gel filtration (Superdex200, GE Healthcare Chicago, IL).

In all cases, the protein concentration was determined using a Bradford Assay (BioRad, Hercules, CA,) with BSA as a standard and protein purity was verified using SDS/PAGE analysis with Coomassie stain. All proteins were flash-frozen and stored at −80°C for longterm storage.

The SNAP25 and Munc13-Halo proteins were fluorescently labeled as described previously (Krishnakumar et al. 2013). Typically, the proteins were incubated with 5X excess dye (AlexaFluor 488, 555 or 660 as noted) for 1h at room temperature (or overnight at 4°C) and the unbound dye was removed using spin columns (Thermofisher, Waltham, MA). The labeling efficiency was estimated using absorbance values (488 or 555 or 660 nm/280 nm) and in all cases, was verified to be >90%.

### Microscale Thermophoresis Interaction Analysis

Halo-tagged Munc13-1 constructs (Munc13L, Munc13_L_^NF^ and MUN) were labeled with AlexaFluor 660 and MST analysis was carried out with 50 nM of fluorescently labeled Munc13-1 constructs in a premium-coated glass capillary with 0.5 mg/ml BSA and 0.05% Tween-20 included in the HEPES buffer (50 mM HEPES, 140 mM KCl, 1 mM TCEP, 1 mM MgCl_2_) to prevent non-specific adsorption to the glass surface. The samples were measured in a Monolith NT.115 (Nanotemper Technologies, San Francisco, CA) and MST curves obtained with titration of SNAP25 (1 nM – 125 μM range) were fitted with a non-linear curve (sigmoidal) analysis to estimate the apparent dissociation constant (K_d_).

### Lipid Bilayer experiments

Liposomes were made using brain physiological composition (73% DOPC, 15% DOPS, 3% PIP2, 2% DAG, 5% DOPE-PEG 2000, 2% NBD-PE) using extrusion method with HEPES buffer (50 mM HEPES, 140 mM KCl, 1 mM TCEP, pH 7.4). Lipid bilayers were created by Mg^2+^(5mM) induced bursting liposomes in ibidi glass-bottom chambers (ibidi GmbH, Germany). The bilayer was extensively washed with HEPES buffer and the fluidity of the lipid bilayer was verified using fluorescence recovery after photo-bleaching (FRAP) using the NBD fluorescence. Consistent with a fluid bilayer, the average diffusion coefficient of the lipid was calculated to be 1.6 ± 0.3 μm^2^/sec (Figure 2 Supplement 1). SNAP25 and Munc13_L_ were added to the pre-washed supported bilayer and incubated for 30 minutes. For the clustering experiments (Figure 2), we used 160 nM SNAP25^WT^ or SNAP25^G4S^ (residue S28C) labeled with AlexaFluor555 with increasing concentrations (50 nm – 2μM) of unlabeled Munc13_L_. FRAP analysis showed that the membrane-bound AlexaFluor555-labeled SNAP25 was fairly mobile, with an average diffusion coefficient of 1.0 ± 0.2 μm^2^/sec (Figure 2 Supplement 1). For the co-localization experiments (Figure 3), we used 160 nM SNAP25^WT^ (residue S28C) labeled with AlexaFluor555 with either 1 μM or 50 nM of Munc13_L_-Halo labeled with AlexaFluor488. For all experiments, the chambers were washed thoroughly (after 30 min incubation) to remove any non-specifically or weakly bound proteins and the samples were imaged on a TIRF microscope (Nikon) with a 63x oil objective.

### FRET Experiment

We used a FRET-based assay to track N-terminal SNARE complex formation. For site-specific labeling with fluorophores, cysteines were introduced into cytoplasmic domains of SNAP25 (residues 20) and VAMP2 (residue 28) and was labeled with thiol-reactive fluorescent probes AlexaFluor488 maleimide and AlexaFluor555 maleimide respectively. 1.5 μM of labeled SNAP25 and unlabeled Syantaxin-1 were pre-mixed with or without 3 μM MUN domain and incubated at 37°C for 10 min. 500 nM of pre-warmed labeled VAMP2 was added to start the reaction and fluorescence at 580 nm was measured in plate reader (Molecular Devices) for 60 min with 15 sec intervals. In control experiments, we left out Syntaxin-1 from the reaction mixture.

### Optical Tweezer experiments

Single molecule force spectroscopy experiments were carried out as described previously (Jiao et al. 2017; Rebane et al. 2016; Shu et al. 2020; Jiao et al. 2018). Briefly, ternary SNARE complexes were assembled by mixing the purified SNARE proteins and incubating at 4°C overnight. Assembled SNARE complexes were purified by binding to Ni-NTA-agarose and cross-linked with DTDP (2,2’-dithiodipyridine disulfide) treated DNA handles (50:1 molar ratio) in PBS buffer (100 mM phosphate buffer, 500 mM NaCl, pH 8.5). Dual-trap optical tweezers and basic protocols for single-molecule experiments have been described in detail elsewhere (Jiao et al. 2017; Rebane et al. 2016). To follow SNARE assembly/disassembly pathways, samples of the 2 DNA handles, one cross-linked with the SNARE complex and the other bound by a streptavidin molecule, were separately bound to anti-digoxigenin antibody coated polystyrene beads. The beads were injected into a microfluidic channel and trapped. The 2 beads are then brought close to allow a single SNARE complex to be tethered between them. All manipulation experiments were carried out in PBS buffer supplemented with the oxygen scavenging system. All single molecules were pulled and relaxed by increasing and decreasing, respectively, the trap separation at a speed of 10 nm/s or held at constant mean forces by keeping the trap separation constant.

## ACKNOWLEDGEMENT

This work was supported by National Institute of Health (NIH) grant DK027044 to JER/SSK and GM131714 to YZ.

---

**Figure 1 Supplement 1.**
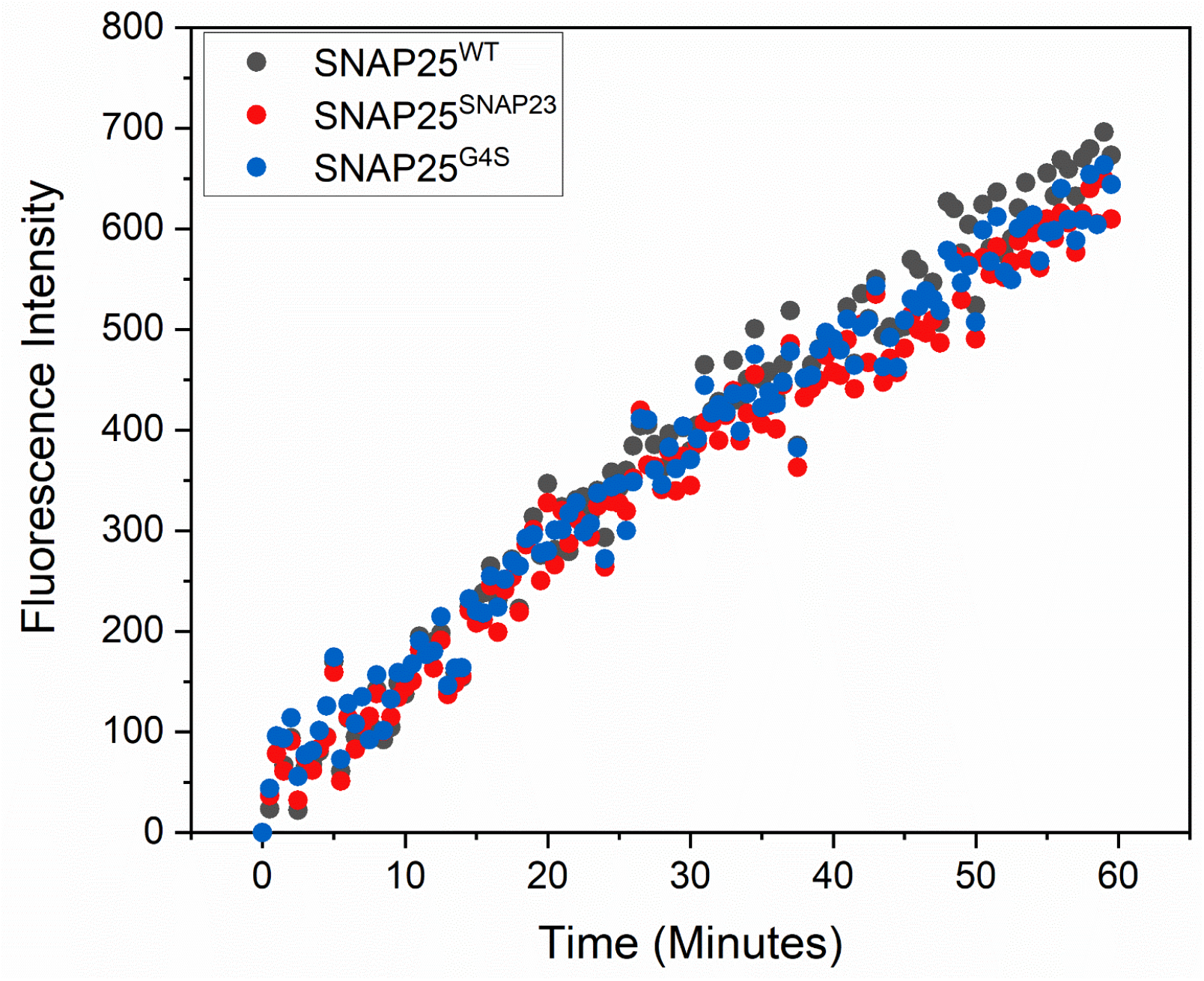
SNAP25 linker domain mutants are functional in a fusion assay. The effect of SNAP25 linker domain mutants (SNAP25^G4S^ and SNAP25^SNAP23^) on SNARE mediated fusion was investigated using the classical lipid mixing assay as described previously (Ji et al. 2010; Weber et al. 1998). Briefly, VAMP2 was reconstituted into donor liposomes (v-liposomes) containing 1.5% NBD and Rhodamine at final 1:160 protein: lipid ratio. The acceptor liposomes were prepared with Syntaxin-1 reconstituted at 1:1600 final ratio. Syntaxin-liposomes were mixed with 2 μM of SNAP25 wild-type or mutant and pre-incubated at 37°C for 5 min, prior to the addition of v-liposomes to start the fusion reaction. Fusion was monitored by the change in the NBD fluorescence (Ex 460 nm, Em 538 nm) resulting from dequenching due to dilution at 37°C using a SpectraMax M5 (Molecular Devices, Sunnyvale, CA) plate reader. The fluorescence signal was normalized against control experiments of v- and t-liposomes mixed together with no SNAP25 added. Both SNAP25^G4S^ and SNAP25^SNAP23^ were found to drive fusion and to levels comparable to the SNAP25^WT^, suggesting that the mutation of the linker region has no effect on ability of SNAP25 to assemble into SNARE complex with Syntaxin-1 and VAMP2.

**Figure 2 Supplement 1.**
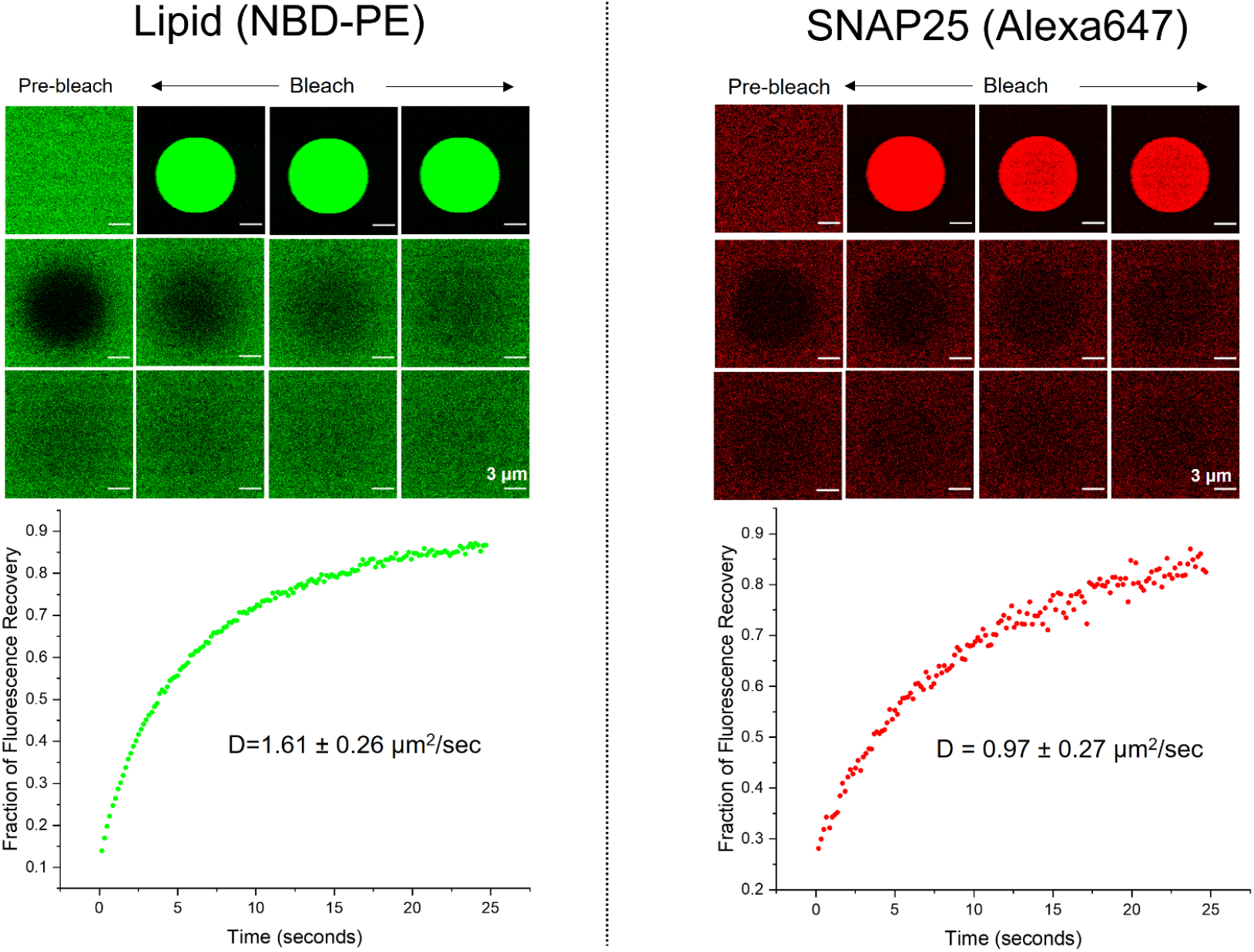
Diffusion of lipid and soluble SNAP25 on supported lipid bilayer. Fluidity of the supported bilayer and mobility of membrane-bound SNAP25 was determined using fluorescence recovery after photo-bleaching (FRAP) experiments. Lipid bilayers containing 2 mol% NBD-PE were bleached with the 488nm laser within a 10 μm ROI and the recovery was measured to determine a diffusion coefficient of 1.6 ± 0.3 μm^2^/sec. Independently, SNAP25-AlexaFluor 647 protein bound to the lipid bilayer surface was bleached using a 647 nm laser within a 10 μm ROI and the recovery was measured to determine a diffusion coefficient of 1 ± 0.3 μm^2^/sec. All experiments were conducted at 37°C using a 40x water objective on a Leica SP8 confocal laser scanning microscope and recovery was tracked for 150 frames at a rate of 147 msec/frame. Representative images for first 10 sec (every 10^th^ frame) are shown. Average from three independent experiments with a minimum of 8 ROIs is shown.

**Figure 3 Supplement 1.**
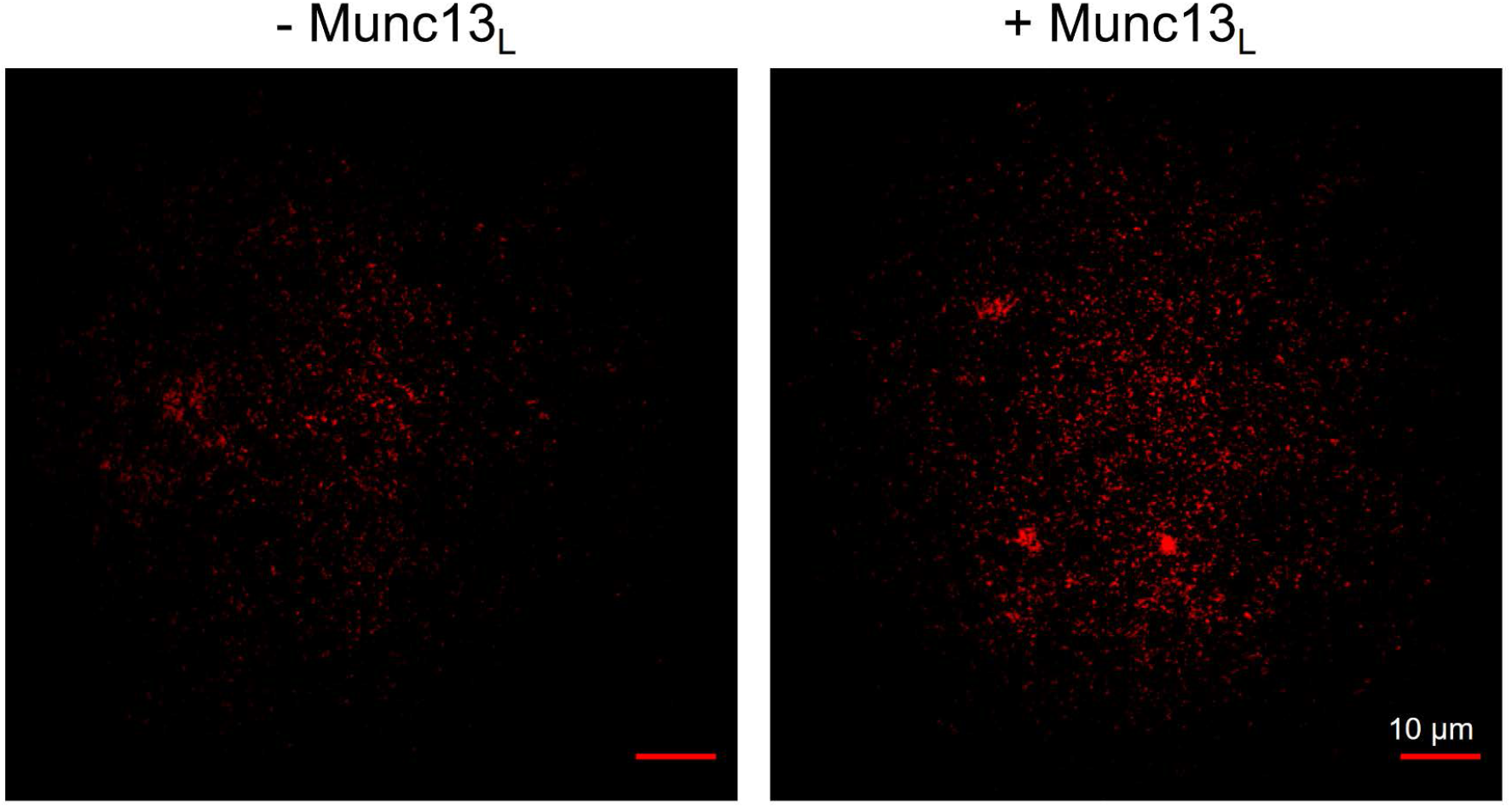
Munc13-1 induces clustering of membrane-anchored SNAP25 molecules. Distribution of AlexaFluor 660 labeled palmitoylated SNAP25^WT^ incorporated into supported lipid bilayer under physiological lipid and buffer composition in the absence or presence of 1 μM Munc13_L_ was visualized using TIRF microscopy. Similar to the membrane-bound soluble SNAP25 molecules (Figure 2), Munc13_L_ induces clustering of the palmitoylated and membrane-anchored SNAP25^WT^. To produce fluorescently labeled palmitoylated SNAP25, SNAP25^WT^ sequence was modified to include a non-natural amino acid (L-azidophenylalanine) at residue 20 using amber stop codon suppression method (Lajoie et al. 2013; Amiram et al. 2015; Chin et al. 2002)and labeled with DBCO-AlexaFluor647 (Jena Biosciences, Jena, Germany) using click chemistry without copper. The labeled SNAP25 protein was palmitoylated by the addition of Palmitoyl-CoA (Sigma Aldrich, St. Louis, MO) at room temperature for 30 minutes and then reconstituted in the liposomes using detergent dilution and dialysis method, and isolated on a discontinuous Nycodenz gradient. Supported lipid bilayers in ibidi chambers were prepared by bursting liposomes using 5 mM MgCl_2_ and 1 μM Munc13_L_ (or buffer) was added to the chambers following extensive wash, incubated for 30 min at 37°C before being washed and imaged using a Nikon TIRF microscope. Representative image from a minimum of three independent trials is shown.

